# A system of reporters for comparative investigation of EJC-independent and EJC-enhanced nonsense-mediated mRNA decay

**DOI:** 10.1101/2023.11.14.567061

**Authors:** Divya Kolakada, Amy E. Campbell, Laura Baquero Galvis, Zhongyou Li, Mlana Lore, Sujatha Jagannathan

## Abstract

Nonsense-mediated mRNA decay (NMD) is a network of pathways that degrades transcripts that undergo premature translation termination. In mammals, NMD can be divided into the exon junction complex (EJC)-enhanced and EJC-independent branches. Fluorescence- and luminescence-based reporters have long been effective tools to investigate NMD, yet existing reporters largely focus on the EJC-enhanced pathway. Here, we present a system of reporters for comparative studies of EJC-independent and EJC-enhanced NMD. This system also enables the study of NMD-associated outcomes such as premature termination codon (PTC) readthrough and truncated protein degradation. These reporters are compatible with fluorescence or luminescence-based readouts via transient transfection or stable integration. Using this reporter system, we show that EJC-enhanced NMD RNA levels are reduced by 2- or 9-fold and protein levels are reduced by 7- or 12-fold compared to EJC-independent NMD, depending on the reporter gene used. Additionally, the extent of readthrough induced by G418 and SMG1i, alone and in combination, varies across NMD substrates. When combined, G418 and SMG1i increase readthrough product levels in an additive manner for EJC-independent reporters, while EJC-enhanced reporters show a synergistic effect. We present these reporters as a valuable toolkit to deepen our understanding of NMD and its associated mechanisms.

## Introduction

RNA quality control mechanisms safeguard cells from aberrant RNAs and their likely deleterious protein products. Nonsense-mediated mRNA decay (NMD) is one such mechanism, that targets aberrant mRNAs that undergo premature translation termination for degradation. NMD substrates can arise from genetic mutations or errors in mRNA synthesis or processing; ∼11.5% of genetic diseases are associated with premature termination codons (PTCs) [1]. NMD also modulates the expression of physiological transcripts, making it integral to biological processes like embryonic development, the integrated stress response, and tissue-specific cell differentiation [2–5]. The central trigger for NMD is an aberrant translation termination event, which is signaled by specific cues [6–10].

The most well-studied cue in mammals and other higher eukaryotes is the presence of a stop codon >50-55 nts upstream of the last exon-exon junction of a transcript [11]. In this case, the sustained presence of an exon junction complex (EJC), an RBP marker of exon-exon junctions, downstream of the terminating ribosome triggers the EJC-enhanced mode of NMD [8, 10, 12]. The other cue relies on the presence of a long 3’ UTR after the normal termination codon (NTC); lacking EJCs, it is EJC-independent [13]. One proposed mechanism for these transcripts involves the long distance between the stop codon and the poly(A) binding protein, which mimics a premature translation termination environment, inducing the EJC-independent mode of NMD [14, 15]. This is the prevalent mode of NMD in *Saccharomyces cerevisiae*, where only 4% of genes contain introns and most EJC factors are not present [16–18]. EJC-independent NMD also occurs in *C. elegans*, *Drosophila*, and mammals, indicating its evolutionary conservation [13, 19, 20].

When a prematurely terminating ribosome is recognized, sustained presence of the EJC or other RBPs downstream serves to recruit the NMD machinery, culminating in the phosphorylation of the central NMD factor UPF1, which triggers degradation of the transcript [21–28]. Compared to EJC-enhanced NMD, less is known about the EJC-independent branch even though approximately 30% of mammalian transcripts have UTRs longer than 1 kb and are potential substrates for this mechanism [29–31]. Moreover, few studies comparatively analyze the EJC-enhanced and EJC-independent modes of NMD. Thus, comparative studies of both pathways are required for a deeper and more complete understanding of this essential RNA quality control mechanism.

NMD does not function like an on/off switch, i.e., not every mRNA with a PTC is degraded [32]. Rather, it exhibits a range of efficiencies, which includes escape from NMD. Factors influencing NMD efficiency include the sequence context, substrate-type (i.e., EJC-enhanced or EJC-independent), or RBPs [13, 30, 33, 34]. For instance, EJC-enhanced substrates are more efficient targets of NMD compared to their EJC-independent counterparts [13, 33]. Further, EJC-independent targets can evade NMD because specific RBPs like PTBP1 bind to their 3’ UTR and prevent UPF1 binding [34]. Other mechanisms of escape from NMD include re-initiation, where a ribosome can re-initiate translation downstream of a PTC, and readthrough, where a ribosome reads through a PTC via incorporation of a non-cognate amino acid [35–38]. Additionally, co- or post-translational protein degradation mechanisms can also target residual NMD-associated truncated protein for degradation, buffering the potential negative effects of NMD escape [39–41]. However, our understanding of these factors and the mechanisms by which they influence NMD efficiency remains incomplete.

Fluorescence- and luminescence-based reporters are powerful tools that enable rapid, sensitive, and quantitative measurements of protein levels, making them compatible with high-throughput approaches like small molecule screening, forward/reverse genetic screens, and massively parallel reporter assays. In 2005, Pailluson *et al.*, published the first fluorescence-based NMD reporter demonstrating that green fluorescent protein (GFP) fluorescence was proportional to RNA levels, thus enabling the study of NMD via flow cytometry, spectrofluorometry, and fluorescence microscopy [42]. Boelz *et al.*, followed with a luminescence-based NMD reporter where the ratio of Renilla to Firefly luciferase reflected RNA levels and responded to small molecule mediated NMD inhibition in a dose-dependent manner, highlighting its utility for small molecule screening [43]. Subsequent reporter systems have incorporated technical modifications such as different fluorescent/luminescent proteins, inducible promoters, and recombinase-mediated stable integration [35, 39, 40, 44–52]. These reporters have been used to demonstrate the variability of NMD efficiency within and between cell types [44, 52]. They have also enabled forward genetic screening, leading to the discovery of new NMD modulators like ICE1, components of the U2 spliceosome, the GIGYF2•EIF4E2 complex, and AKT signaling [46, 48, 50, 51, 53]. Moreover, they have allowed for further study of PTC-associated readthrough, and the discovery of truncated protein degradation associated with NMD [35, 39, 40, 47, 52].

Existing fluorescence- and luminescence-based NMD reporters largely focus EJC-enhanced NMD while only a small subset examine EJC-independent NMD [35, 39, 42–46, 48–52]. There are two reporter sets that target both pathways, however they were designed to study NMD-associated truncated protein degradation, necessitating modifications to assess other aspects of NMD [40, 41]. The predominant focus on only one mode of NMD makes it difficult to know whether discoveries from one reporter can apply to both mechanisms, highlighting the need for a reporter system for comparative analysis. To address these limitations, we introduce a reporter system designed to maximize the number of NMD-influencing variables that can be tested in parallel. These reporters enable the comparative study of EJC-enhanced and EJC-independent NMD using either fluorescent or luminescent reporter genes. Our reporters are compatible with transient transfections or stable integrations and can also be used for high throughput assays, genetic screens, or massively parallel reporter assays. Finally, our reporters can investigate at least two NMD-associated phenomena: PTC readthrough and truncated protein degradation. We present this system of reporters as a resource to the community for advancing our understanding of NMD and its underlying mechanisms.

## Materials and Methods

### Plasmid construction

Fluorescent reporters: RFP (TurboFP365), GFP (TagGFP2) with and without a GAPDH intron, and BFP (mAzurite) were synthesized as gene fragments (Twist Biosciences). These fragments were cloned into a Lox/LoxP backbone (obtained courtesy of the Taliaferro lab) via InFusion cloning (Takara) to assemble the NMD+ EJC-independent and EJC-enhanced reporters. The NMD– reporter was created by restriction digesting the EJC-independent backbone with EcoRI and MfeI, filling the sticky ends with Klenow and ligating the blunt ends together to remove the 3’ UTR contributed by GFP. Luminescent reporters: The fragment containing RFP, CMV promoter, and BFP sequences was removed by restriction digestion and replaced with a synthesized gene fragment containing Firefly luciferase, CMV promoter, and Renilla luciferase sequences. All plasmids were confirmed via whole plasmid sequencing. The Firefly/Renilla luciferase gene fragment was ordered as a single fragment from Twist Biosciences. The 5’UTR synthetic intron fragment was synthesized (Twist Biosciences) and cloned into the AgeI restriction site in all reporter plasmids via InFusion (Takara). These plasmids are available through Addgene (Addgene ID#s available upon publication).

### Cell culture, transfections, and genomic integration

HEK293T cells were cultured in Dulbecco’s DMEM with 10% EqualFetal. Cells were transfected in 6-well plates at a 40% confluency with 2500 ng of plasmid DNA using Lipofectamine 2000 (Thermo), as per manufacturer’s instructions.

Transfections for fluorescent and luminescent reporters were performed differently since fluorescent proteins have longer maturation rates than luciferase proteins [54]. For the fluorescent reporters, cells were split 24 hours after transfection equally into two new 6-wells. Forty-eight hours after transfection, one set of cells were harvested for RNA and the other set for flow cytometry. For the luminescent reporters, cells were harvested 24 hours after transfection in PBS with 60% of each well harvested for RNA and 30% pelleted and frozen for a luciferase assays.

Due to the difference in protein maturation rates, SMG1i treatment was also slightly different for each set of reporters. Fluorescent reporters were treated with 0.5 μM of SMG1i in DMSO for 20 hours before harvest. Luminescent reporters were treated with 1 μM of SMG1i in DMSO for 4 hours before harvest. The negative control was DMSO without any added drug. For the experiments with fluorescent reporters and G418, G418 was added to cells transfected with reporters 20h before harvest at a final concentration of 2 mg/mL.

To genomically integrate the reporters into HEK293T cells containing a Lox2272 and LoxP enclosing a blasticidin cassette (courtesy Taliaferro Lab, CU Anschutz), these cells were co-transfected with 97.5% NMD reporter and 2.5% Cre-plasmid. Twenty-four hours after transfection, the cells were provided with fresh media. Puromycin selection began 48 hours after transfection (puromycin concentration: 2 μg/mL). Puromycin selection lasted between 1-2 weeks, during which the media was replaced with fresh media supplemented with puromycin every 1-2 days. Cell lines obtained at the end were maintained in media containing puromycin.

### RNA harvest, cDNA synthesis, and RT-qPCR

RNA was harvested from cells using the RNeasy Mini Kit (Qiagen) lysis buffer. The RNA extractions were subsequently performed as per manufacturer’s instructions. 5 ug of extracted RNA was subjected to a DNAse digest using Turbo DNAse (Thermo), as per manufacturer’s instructions. To inactivate the enzyme, EDTA was added at a final concentration of 15 mM and samples were incubated at 75°C for 10 minutes. cDNA was created from 200 ng of DNase digested RNA using Superscript III First Strand Synthesis kit (Invitrogen) as per manufacturer’s instructions. A no-RT sample was included as a control to make sure there was no gDNA contamination.

The cDNA was then diluted 1:4 and 2 μL were used per 5 μL qPCR reaction. qPCR was performed using iTaq Universal SYBR Green Supermix (Bio-Rad) and primers specific for BFP and RFP or Firefly luciferase and Renilla luciferase, depending on the reporters used. Primers were used at a final concentration of 0.25 μM. Their sequences were as follows:

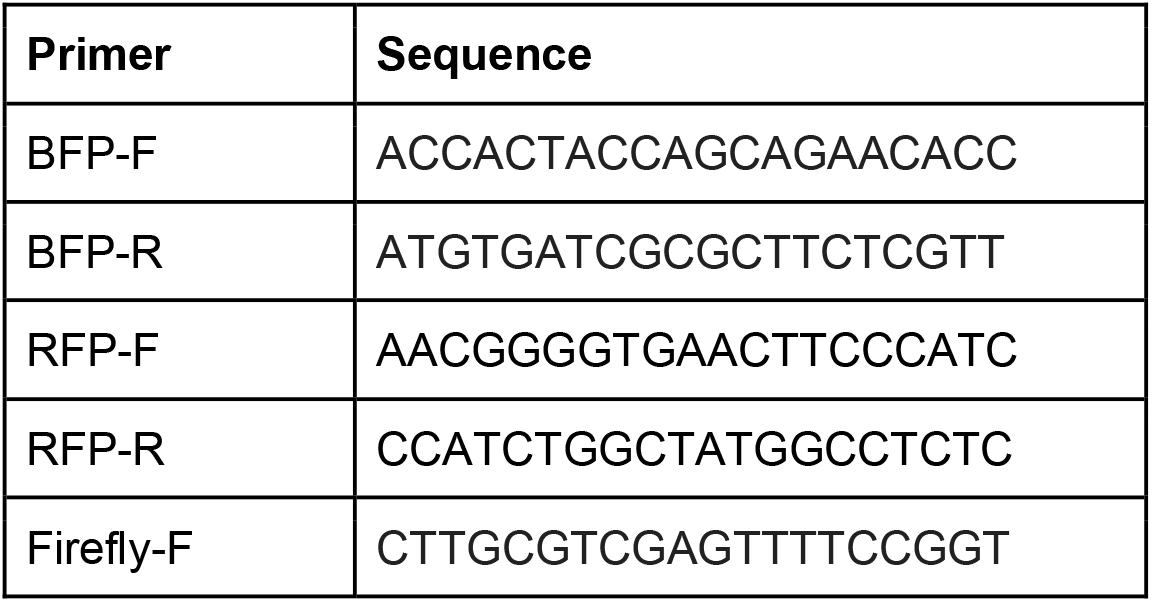
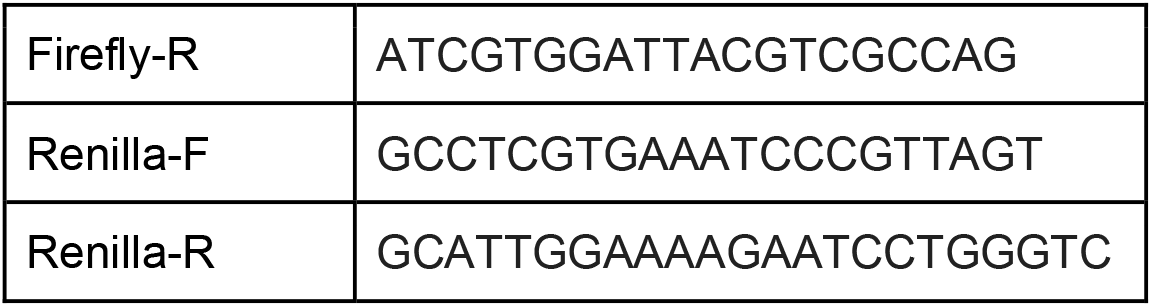

Reactions were set up in triplicate per sample and plated in 384-well plates. The plates were then run on the OPUS Bio-Rad qPCR machine using the 2-step Amp + melt curve protocol. The delta Ct method was used to quantify RNA levels: RFP was normalized to BFP, and Firefly luciferase was normalized to Renilla luciferase. For the transient transfections and genomic integrations, the mean RNA levels of 3 different transfections or 3 technical replicates were plotted, respectively. Error was calculated using standard error of the mean.

### Flow Cytometry

Cells were washed with 1 mL of PBS and pelleted at 400 *xg* for 5 minutes. The cells were then incubated in a 1:1000 dilution Ghost Dye Red 780 viability dye in PBS for 20 minutes at 4°C, in the dark. The cells were pelleted at 400 *xg* for 5 minutes and resuspended in 400 μL of PBS, before being analyzed by the Agilent Novocyte Penteon flow cytometer at the CU Cancer Center Flow Cytometry Core. Wavelengths used were RFP (excitation: 588, emission: 635), GFP (excitation: 483, emission: 506), and BFP (excitation: 384, emission: 450), and Ghost dye (excitation:633, emission: 780). Data were collected from 100,000 cells and RFP expression was analyzed from live cells that were positive for BFP.

### Luciferase Assays

Luciferase assays were performed using the Dual-Luciferase Reporter Assay System (Promega). Frozen pellets were resuspended in 300 μL of 1X Passive Lysis Buffer (PLB) and passively lysed on a rocker for 15 minutes. Following lysis, pellets were vortexed to make sure they were lysed thoroughly, 30 seconds per sample. A portion of each sample of lysed cells was diluted 1:16 in 1X PLB. Twenty μL of this dilution was pipetted into a 96-well plate, in triplicate, per sample. The LAR II and Stop & Glo reagents were prepared as per manufacturer’s instructions, 100 μL per sample, each. Luciferase assay was performed using the Glomax Navigator as per the Dual Luciferase Assay Protocol.

### Western blotting

Cells were harvested in RIPA buffer supplemented with complete protease inhibitor (Roche). Protein was run on NuPAGE Bis-Tris precast polyacrylamide gels (Thermo Fisher Scientific) alongside PageRuler Plus Prestained Protein Ladder (Thermo Fisher Scientific) and transferred to Odyssey nitrocellulose membrane (LI-COR Biosciences). Membranes were blocked in Intercept (PBS) Blocking Buffer (LI-COR Biosciences) before overnight incubation at 4°C with primary antibodies diluted in Blocking Buffer containing 0.2% Tween 20. Membranes were incubated with IRDye-conjugated secondary antibodies (LI-COR Biosciences) for 1 h and fluorescent signal visualized using a Sapphire Biomolecular Imager (Azure Biosystems) and Sapphire Capture software (Azure Biosystems). When appropriate, membranes were stripped with Restore Western Blot Stripping Buffer (Thermo Fisher Scientific) before being re-probed.

### Reagents

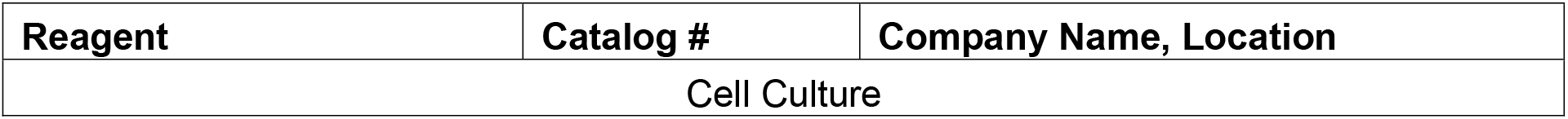
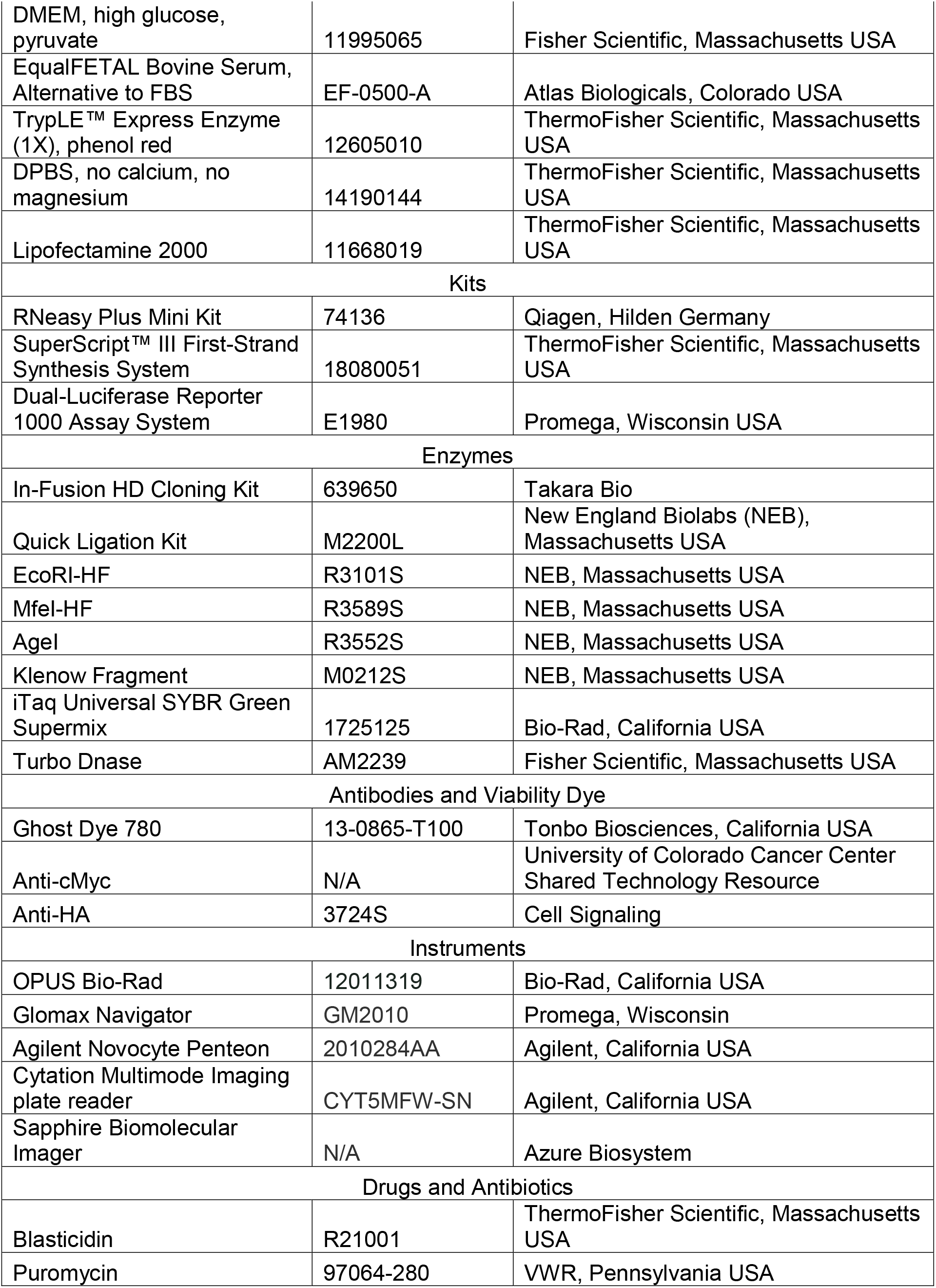
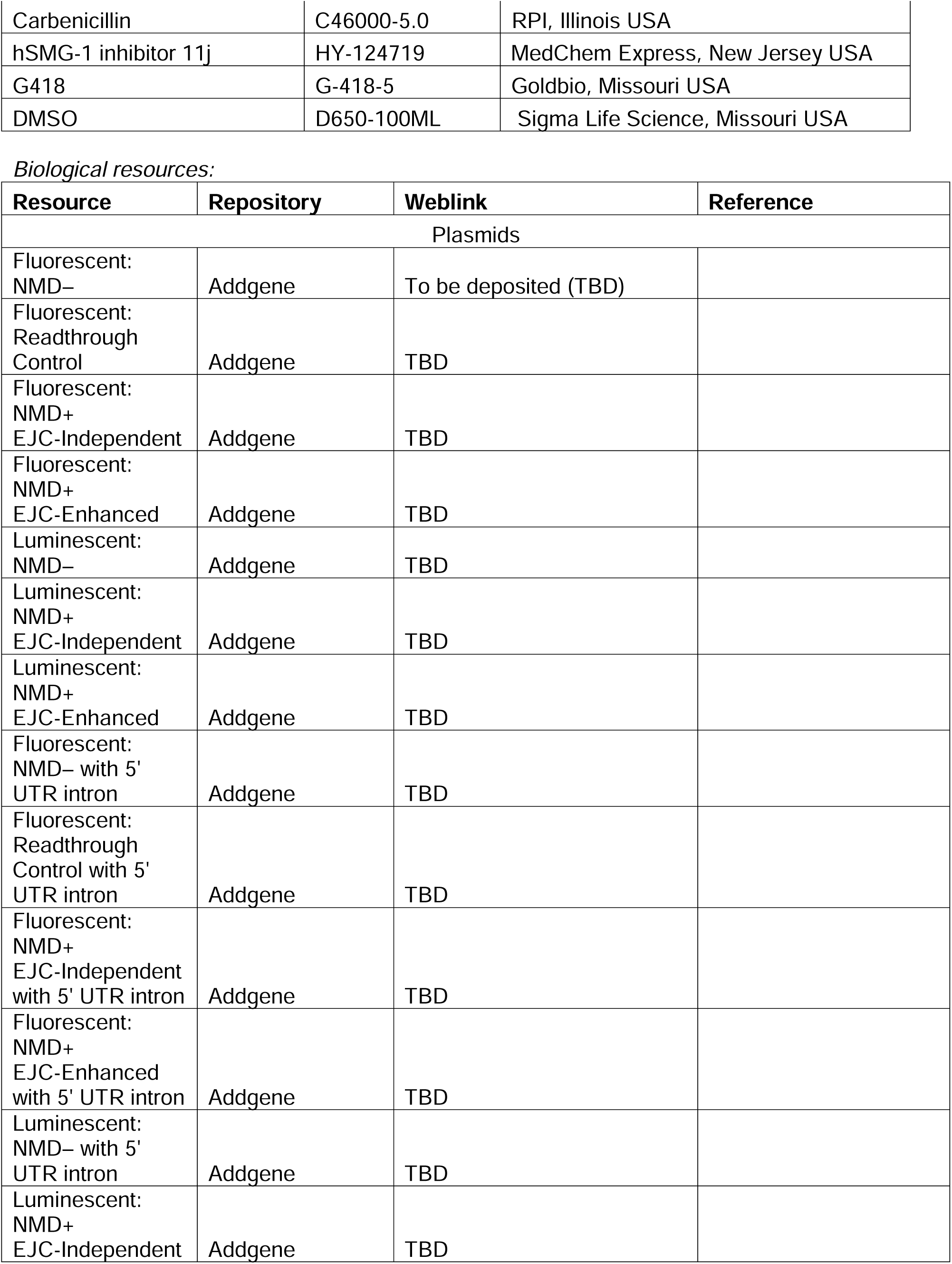
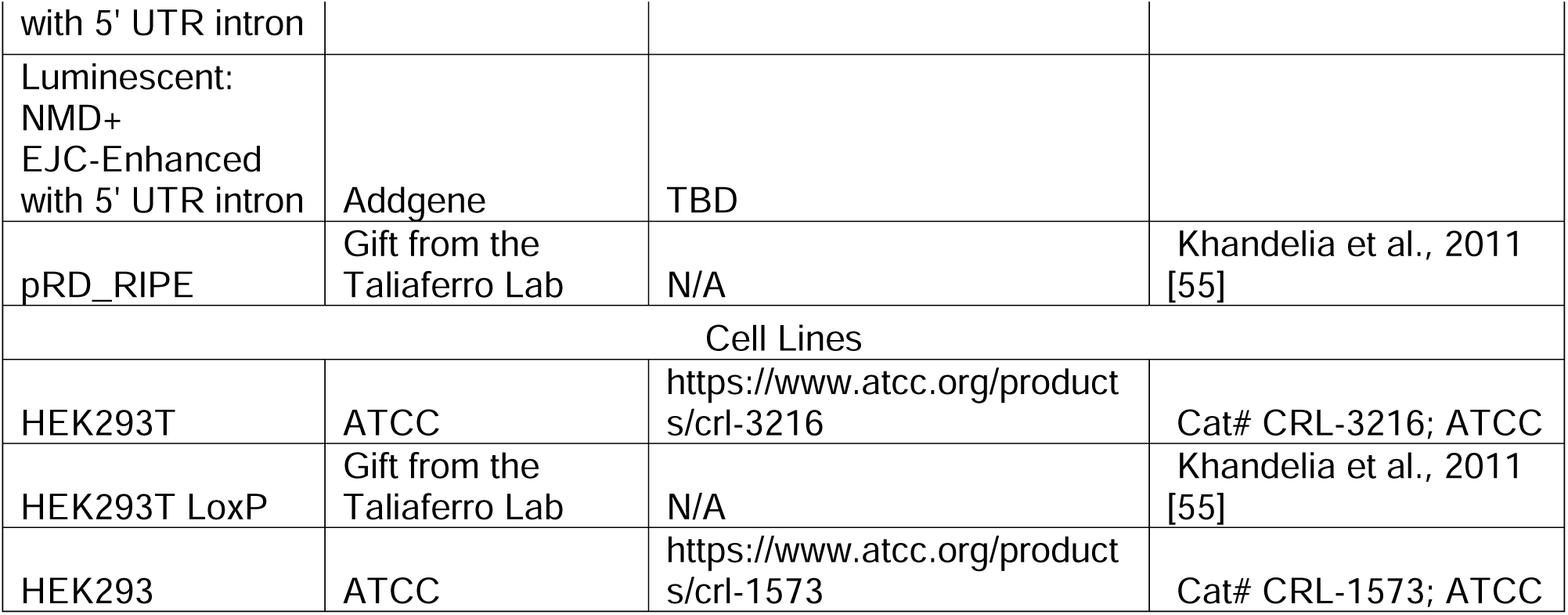

### Statistical Analyses

All experiments were performed with a sample size of n=3. For transient transfections, each replicate represents an independent transfection. For stably integrated cells, the same cell line was plated in 3 different wells, from which RNA and protein were harvested. Statistical significance between various populations was calculated using a student’s t-test using the ggpubr R package and p-values were two-sided (*p<0.05, **p<0.005, ***p<0.0005) [56].

### Data Availability/Sequence Data Resources

N/A

### Data Availability/Novel Programs, Software, Algorithms

N/A

### Websites/Data Base Referencing

N/A

## Results and Discussion

### Design of the NMD reporter system for comparative EJC-independent and EJC-enhanced studies

To enable robust comparative studies of NMD mechanisms, we developed two versions of NMD reporters: fluorescence-based reporters to facilitate genetic screens for NMD modulators, and luminescence-based reporters for applications that require more sensitivity to capture variation in NMD. Used in parallel, these reporters also allow the study of transcript-dependent variability in NMD efficiency and associated phenomena such as PTC-readthrough and truncated protein degradation.

Each NMD reporter version contained an EJC-independent and an EJC-enhanced NMD positive (NMD+) reporter, as well as an NMD negative (NMD–) control (**Fig. 1A**). The NMD+ fluorescent reporters consisted of a red fluorescent protein (RFP) (**Fig. 1A**, R2), followed by a GFP open reading frame (**Fig. 1A**, R3), where a stop codon after RFP served as the PTC. The 3’ UTR formed by the intron-less GFP or the intron-containing GFP targeted the transcripts to the EJC-independent or -enhanced modes of NMD, respectively. For the NMD+ luminescent reporters, RFP was replaced by Firefly luciferase (**Fig. 1A**, R2). Finally, the ribosomal skipping signal T2A ensured independent production of the R2 and R3 proteins. Note that R3 has no start codon, which prevents reinitiation of R3 translation. The NMD– reporter contained all the elements of the NMD+ reporters except the T2A signal and the GFP long 3’ UTR, thus, the stop after RFP became a normal stop and the transcript was not a target for NMD (**Fig. 1A**). Expression of each of these reporters was driven by a bi-directional CMV promoter, which also drove expression of a transfection control (**Fig. 1A**, R1; blue fluorescent protein (BFP) or Renilla luciferase, respectively, for the fluorescence- and luminescence-based reporters).

**Fig. 1:**
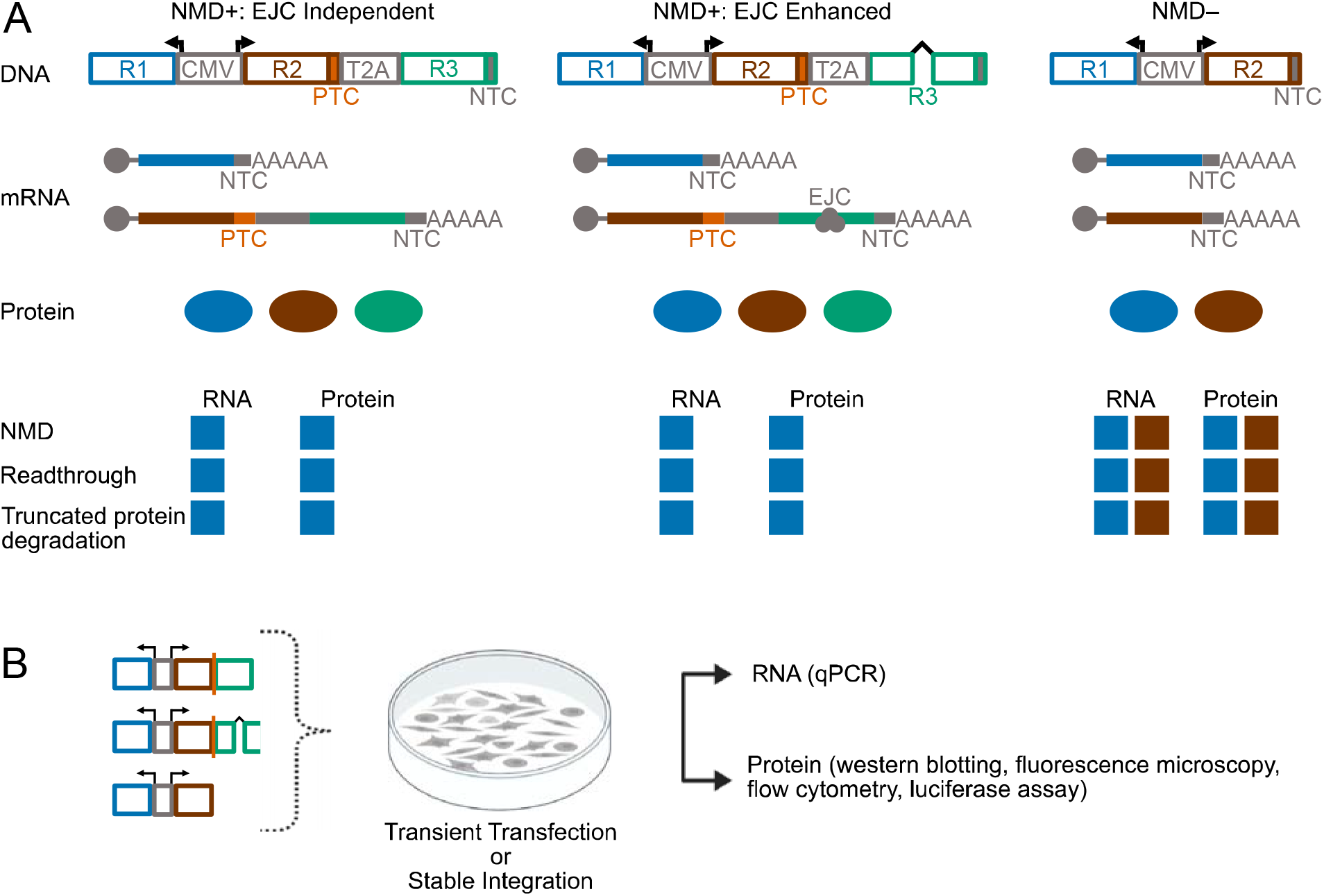
The EJC-independent and EJC-enhanced reporter system design. (A) Top: The NMD+: EJC-independent, NMD+: EJC-enhanced, and NMD– reporter designs, as well as the theoretical mRNAs and proteins that the plasmid reporters can produce. Bottom: The theoretical RNA and protein levels expected from the reporters for NMD and its associated phenomena. (B) Experimental workflow for these reporters as well as possible assays to measure RNA and protein levels. R1 and R2 are either BFP and RFP (respectively), or Renilla and firefly luciferase (respectively) depending on fluorescent or luminescent readouts. R3 is always GFP.

While compatible with transient transfections, these reporters are also flanked by Lox2272 and LoxP sites, to enable recombinase-mediated stable integration into cells with the corresponding Lox sites. Further, the R2 and R3 proteins had epitope tags on their N- and/or C-terminal ends to monitor their translation products. R2 has a N-terminal Myc tag; a C-terminal 3xFLAG tag is positioned after the stop codon of R2 to detect stop codon readthrough; R3 has an N-terminal V5 tag and a C-terminal HA tag. These epitope tags were designed to facilitate sensitive characterization as well as affinity isolation of the various protein products from the NMD reporters to further investigate NMD-associated outcomes (**Supplementary Fig. 1A**; example western blot in **Supplementary Fig. 1B**). In addition to these features, the reporters also have convenient restriction sites that enable straightforward modification of any portion of the reporter cassette (**Supplementary Fig. 1A**).

Each reporter produces two transcripts: the transfection control R1 and the NMD +/– reporter transcript comprising R2 and R3. The transcripts theoretically produce three separate proteins depending on whether they support stop codon readthrough and the extent to which they undergo NMD or NMD-associated protein degradation (middle, **Fig. 1A**). For all reporters, R1 RNA and protein levels would remain consistent. If targeted by NMD, the R2 RNA and protein levels for NMD+ reporters would be less than the NMD– control, with the EJC-enhanced reporters undergoing stronger reduction compared to the EJC-independent reporters. If there was readthrough, then RNA and protein levels of R2 would be comparable to transcripts undergoing NMD but R3 protein expression would also be detected. If there was NMD-associated truncated protein degradation, then the RNA levels of R2 would be comparable to those of the NMD scenario but the protein levels of R2 for the NMD+ reporters would see a stronger reduction compared to the RNA levels (bottom, **Fig. 1A**). Upon transfection or stable integration of these reporters into cells, RNA levels could be measured via RT-qPCR and protein levels by western blotting, fluorescence microscopy, flow cytometry, and luciferase assays (**Fig. 1B**).

### The EJC-independent and EJC-enhanced reporter system enables quantitative investigation of variable NMD efficiency

To determine whether the reporters were targeted by NMD, we transiently transfected them into HEK293T cells and treated with the SMG1 kinase inhibitor (SMG1i) to inhibit NMD[57]. Reporter RNA levels were measured by RT-qPCR, with RFP levels normalized to BFP and Firefly levels normalized to Renilla. As expected, the EJC-independent NMD+ reporter RNA levels showed a statistically significant decrease compared to the NMD– RNA levels (fluorescent: ∼9-fold, luminescent: ∼6-fold; **Fig. 2A-B**). The EJC-enhanced NMD+ reporters also followed the same trend with RNA levels lower than NMD– levels (fluorescent: ∼18-fold decrease, luminescent: ∼55-fold decrease; **Fig. 2A-B**). The decrease in RNA levels of EJC-enhanced reporters was stronger by ∼2 fold (fluorescent) or ∼9 fold (luminescent) compared to EJC-independent NMD+ substrates (**Fig. 2A-B**). Inhibition with SMG1i increased EJC-independent NMD+ reporter RNA levels (fluorescent: ∼2 fold, luminescent: ∼2 fold), as well as EJC-enhanced levels (fluorescent: ∼5-fold, luminescent: ∼15-fold). As expected, SMG1i treatment did not affect NMD– RNA levels (**Fig. 2A-B**). Taken together, these data show that the EJC-independent and EJC-enhanced reporter transcripts are targeted by NMD.

**Fig. 2:**
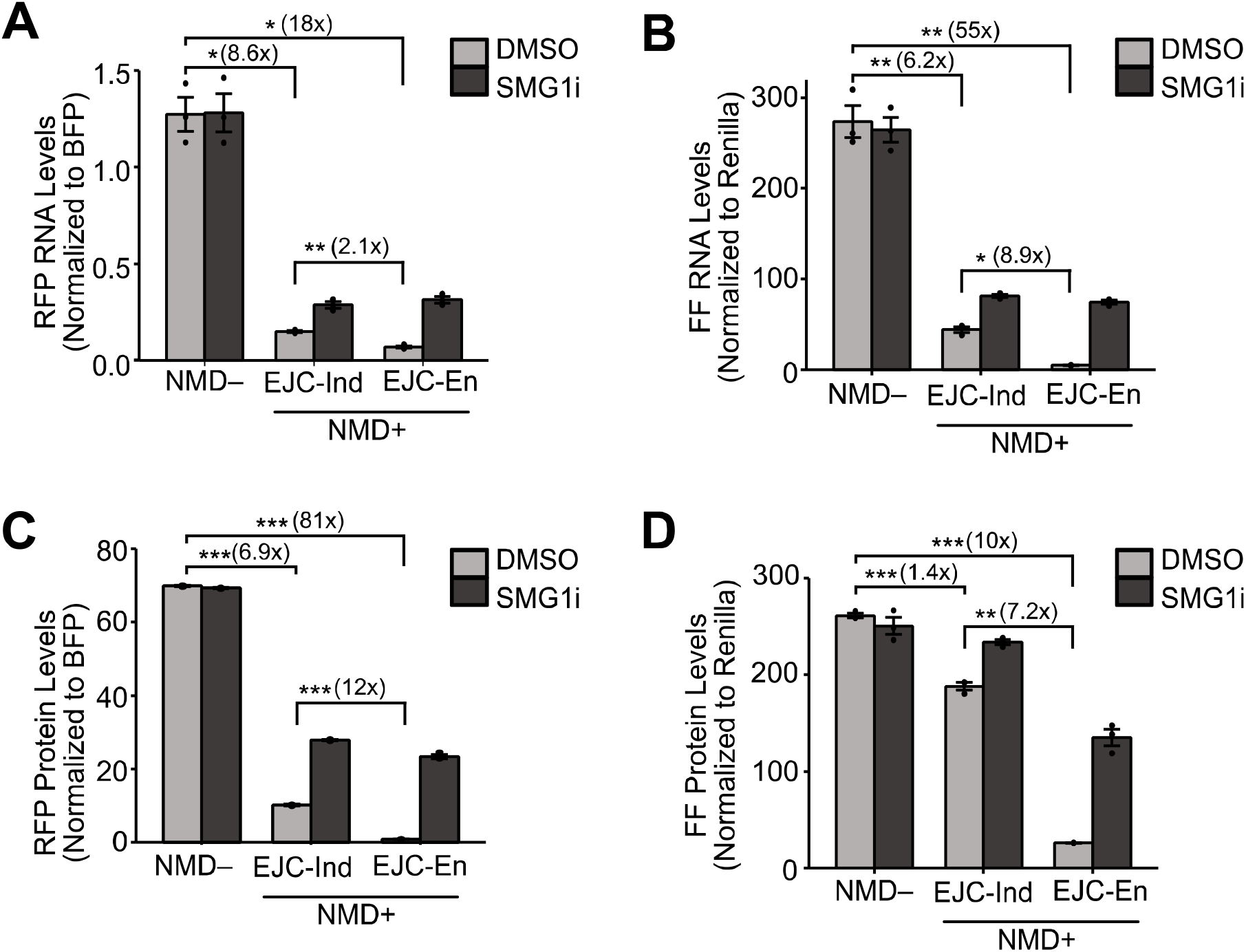
The EJC-independent and EJC-enhanced reporter system enables quantitative investigation of variable NMD efficiency. RNA levels for fluorescent reporters (A) and luminescent reporters (B) as measured by RT-qPCR. Protein levels for fluorescent reporters (C) measured by flow cytometry and plotted as a bar graph using the median RFP intensities. Protein levels for luminescent reporters (D) as measured by luciferase assays. FF refers to Firefly luciferase. Transfections were performed 3 times and the mean values of RNA and protein levels were plotted in the bar graphs. Dots overlaid on each bar indicate the individual data points for that sample. Error bars reflect the standard error of the mean. T-tests were conducted to determine statistical significance (*p<0.05, **p<0.005, ***p<0.0005).

Reporter protein levels were measured via flow cytometry or luciferase assays. For the fluorescent reporters, live cells were gated, and from these only BFP-positive cells were analyzed for RFP expression (gating strategy **Supplementary Fig. 2A-D**). The mean of the median intensities of RFP was then normalized to BFP. For the luminescent reporters, Firefly luminescence was normalized to Renilla. The EJC-independent reporter protein levels decreased compared to NMD– levels (fluorescent: ∼7, luminescent: ∼1.4-fold; **Fig. 2C-D**). The EJC-enhanced reporters followed the same pattern with a much stronger decrease (fluorescent: ∼81-fold, luminescent: ∼10-fold; **Fig. 2C-D**). The extent of reduction of the EJC-enhanced reporters was stronger by ∼12-fold (fluorescent) or ∼7-fold (luminescent) compared to the respective EJC-independent reporters (**Fig. 2C-D**). SMG1i treatment increased EJC-independent NMD+ reporter protein levels (fluorescent: ∼3 fold, luminescent: ∼1.2 fold), as well as EJC-enhanced levels (fluorescent: ∼27-fold, luminescent: ∼5-fold). SMG1i treatment did not affect NMD– protein levels (**Fig. 2C-D**). Overall, protein levels largely reflected RNA levels, suggesting that the reporters are faithful proxies of NMD activity in the cell.

The RNA and protein level data confirmed that our reporters were targeted for degradation by NMD. NMD affected the fluorescence- and luminescence-based reporters to different degrees and this difference was influenced by the mode of NMD that acted on the reporter. Previous research has suggested that protein levels from NMD substrates are typically reduced more strongly than RNA levels, suggesting that protein degradation mechanisms work in conjunction with NMD to eliminate truncated protein [39, 40]. In this study, this pattern only applied to the EJC-enhanced fluorescent reporter. With this reporter, the RNA decreased by ∼18-fold and the protein decreased by ∼81-fold compared to respective NMD– levels. Further, SMG1i treatment increased the RNA and protein to levels comparable to that of the EJC-independent fluorescent reporter when treated with SMG1i. Taken together, these data suggest that mechanisms that may alter the protein output from an NMD target are likely influenced by the precise mode of NMD and/or the type of protein substrate. Such mechanisms may include differential export rates of the transcripts, altered translation efficiency, as well as co- and post-translational protein degradation. Further, despite all reporter RNA and protein levels increasing upon SMG1i treatment, none of the NMD+ reporters reached NMD– levels except for the luminescent EJC-independent reporter, which is closest to NMD– levels. This could be due to the different sequence and structure of the NMD– reporter, which lacks the T2A sequence and R3 and thus, has a short 3’ UTR, potentially altering stability, export kinetics, or translation efficiency.

One aspect to note here is that these NMD– and EJC-independent reporters do not contain an intron anywhere in the transcript. Given that splicing is known to influence RNA export and translation, we generated another set of reporters where a synthetic intron was inserted in the 5’ UTR upstream of R2. These reporters behaved almost identically to the original set of reporters with the EJC-enhanced reporters showing the greatest NMD sensitivity followed by EJC-independent and the controls remained insensitive to NMD (**Supplementary Fig. 3 A-B**). Hence, either of our reporter sets can be utilized to study NMD activity depending on the investigation. Altogether, these reporters enabled quantitative comparisons between the EJC-independent and EJC-enhanced modes of NMD.

### Genomically integrated NMD reporters show less cell-to-cell variability in NMD activity than transiently expressed reporters

As transient transfection is known to impact NMD efficiency, stable integration of reporters is often preferred [52, 58]. Stable integration has other advantages such as making the reporter system amenable to genetic screens and massively parallel reporter assays. To test whether the expression patterns of our NMD reporters remained consistent with stable integration, we used the high-efficiency and low-background recombinant-mediated cassette exchange (HILO-RMCE) approach described by Khandelia et al., to yield HEK293T cell lines that stably expressed a single copy of our fluorescent reporters (**Fig. 3A**) [55].

**Fig. 3:**
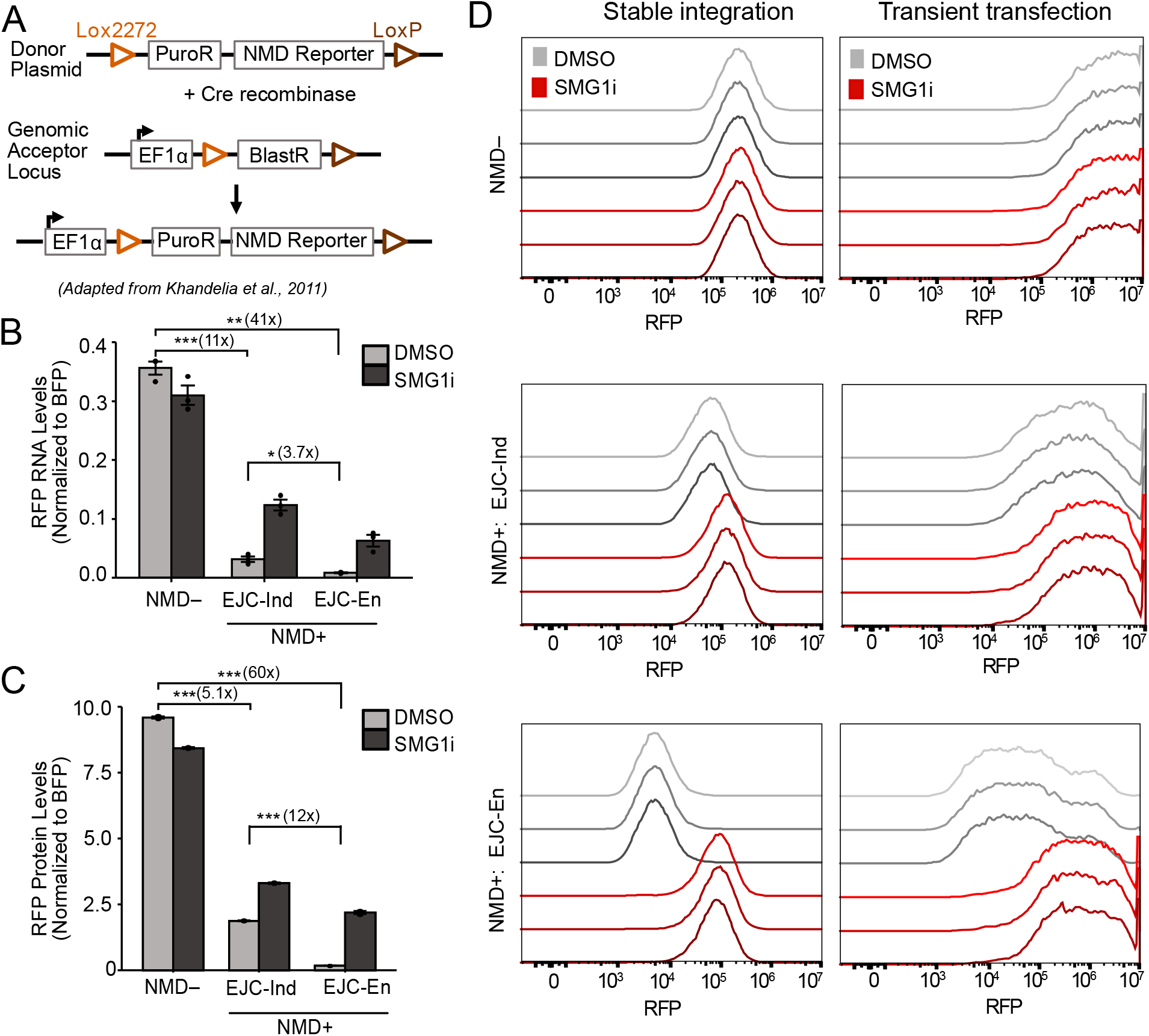
Genomically integrated NMD reporters show less cell-to-cell variability in NMD activity than transiently expressed reporters. (A) Schematic of stable integration using the HILO-RMCE method described by Khandelia et al. [55] (B) RNA levels for stably integrated fluorescent reporters as measured by RT-qPCR. (C) Protein levels for stably integrated fluorescent reporters measured by flow cytometry and plotted as a bar graph using the median RFP intensities. Data obtained from 3 technical replicates of stably integrated cell lines and the mean values of RNA and protein levels were plotted in bar graphs. Dots overlaid on each bar indicate the individual data points for that sample. Error bars reflect the standard error of the mean. T-tests were conducted to determine statistical significance (*p<0.05, **p<0.005, ***p<0.0005). (D) Flow cytometry histograms showing the RFP distribution of BFP+ cells for NMD+: EJC-independent, NMD+: EJC-enhanced, and NMD– reporters, when transiently transfected or stably integrated. The three shades of gray (DMSO treatment) or red (SMG1i treatment) represent different replicates.

The RNA and protein level patterns in stable cells were largely consistent with the results obtained by transient transfection. The EJC-independent and EJC-enhanced reporters showed a decrease in RNA levels (∼11 versus ∼41-fold) compared to NMD– control (**Fig. 3B**). This decrease in EJC-independent RNA was comparable to that observed by transient transfection, but the decrease in EJC-enhanced RNA was more pronounced. SMG1i treatment resulted in an increase in RNA levels for the EJC-independent and EJC-enhanced reporters (∼4 versus ∼7-fold), which were similar in magnitude to those seen during transient transfection (**Fig. 3B**). Protein levels followed the same pattern as RNA levels with a ∼5- and ∼60-fold reduction for EJC-independent and EJC-enhanced reporters, respectively. Upon SMG1 inhibition, protein levels increased by ∼2- versus ∼14-fold for the EJC-independent versus EJC-enhanced reporters (**Fig. 3C**). As expected, SMG1i treatment did not affect stably integrated NMD– RNA or protein levels (**Figs. 3B-C**).

The reduction in EJC-independent protein levels and their increase upon SMG1 inhibition were similar between transient transfection and stable integration. In contrast, the decrease in protein levels and the increase upon SMG1 inhibition for the EJC-enhanced reporter were less pronounced in stable integration compared to transient transfection. Nevertheless, the overall trend remained consistent, as did the enhanced reduction of protein levels compared to RNA levels for the EJC-enhanced reporter. Interestingly, comparing the histograms for stable integration and transient transfection revealed that NMD in a cell population is less variable for stable integration compared to transient transfection, indicating that stable integration of reporters is a more reliable method for determining NMD activity (**Fig. 3D**).

### EJC-independent and EJC-enhanced reporters respond differentially when treated with readthrough-inducing drugs

Readthrough at a PTC can allow a transcript to escape NMD and prevent loss of function of the gene containing a nonsense mutation [29, 59]. Hence, understanding the mechanisms underlying readthrough is essential to developing therapies for NMD-associated diseases [59]. To determine whether our reporter system could be used to study readthrough, we first designed a readthrough control. This NMD– reporter was identical to the EJC-independent NMD+ reporter, except that the PTC was replaced with a Tyr codon (TAC) (**Fig. 4A**, top). Thus, RFP and GFP were both part of the transcript open reading frame but independently expressed due to the T2A signal, and GFP protein expression from this transcript could be used as a positive control for readthrough. The readthrough control and the NMD+ reporters were transiently transfected into HEK293 cells and treated with DMSO, the readthrough-inducing drug G418, NMD inhibitor SMG1i, or a combination of SMG1i and G418 [60]. HEK293 cells were utilized for this set of experiments instead of HEK293T cells since the latter are resistant to G418. BFP, RFP, and GFP levels were measured via flow cytometry. Readthrough was measured by the percentage of cells that were GFP positive, henceforth referred to as GFP-positive cells (**Fig. 4A-B**, bottom, black arrows), as well as by the fluorescence intensities of GFP from the BFP positive cell population, referred to as GFP protein levels (**Fig. 4C**).

**Fig. 4:**
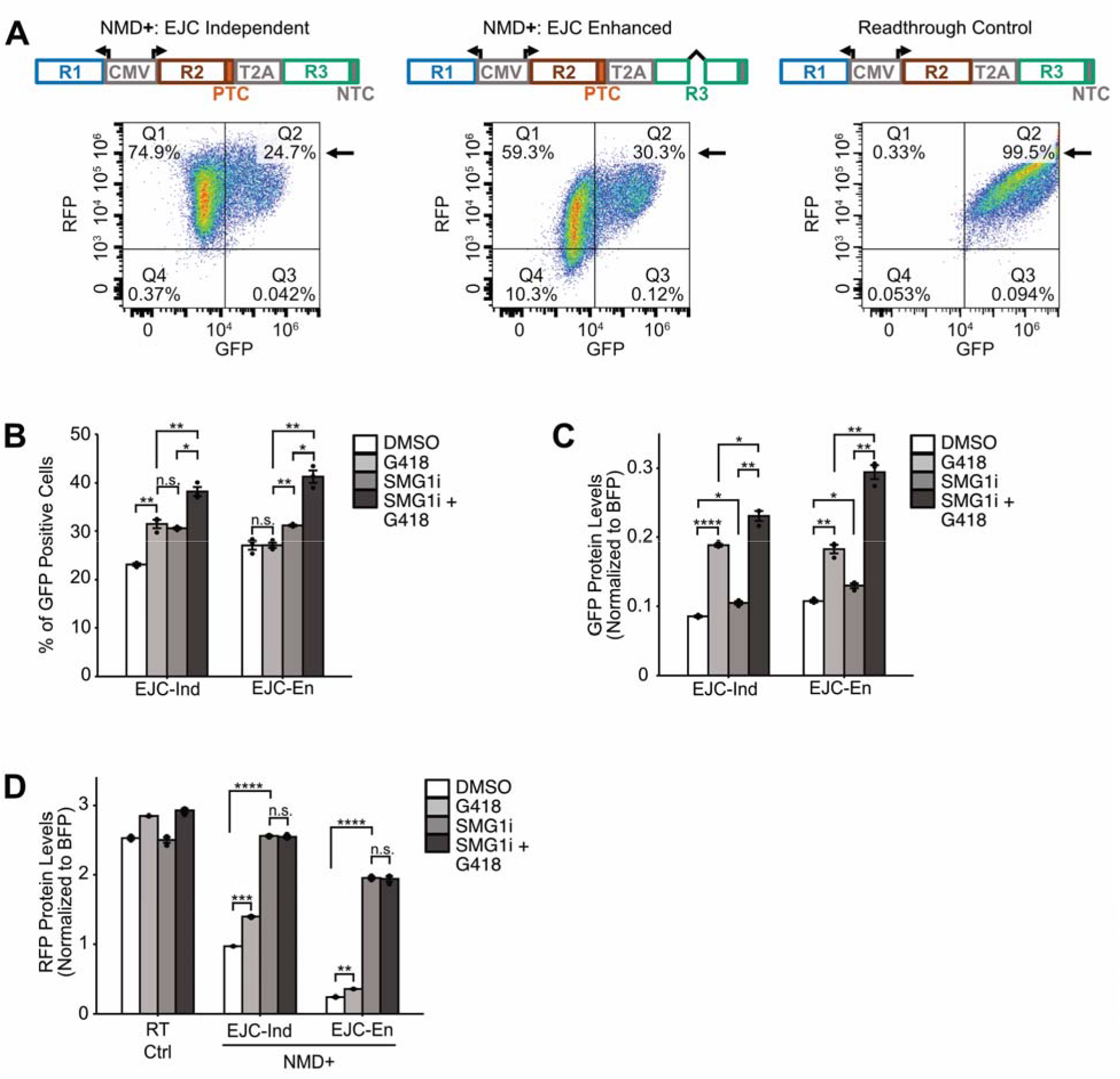
EJC-independent and EJC-enhanced reporters respond differentially when treated with readthrough inducing drugs. (A) Top: schematic of NMD+ and readthrough control. R1 is BFP, R2 is RFP, and R3 is GFP. Bottom: Representative flow diagrams for each reporter. The black arrow is pointing to the quadrant of interest which shows the percentage of cells that are GFP positive. (B) Percentage of GFP positive cells for NMD positive reporters when treated with DMSO, G418, SMG1i, and SMG1i+G418. (C) GFP protein levels normalized to BFP for each of the reporters when treated with DMSO, G418, SMG1i, and SMG1i+G418. (D) RFP protein levels normalized to BFP for each of the reporters when treated with DMSO, G418, SMG1i, and SMG1i+G418. Transfections were performed 3 times and the mean values plotted in the bar graphs. Dots overlaid on each bar indicate the individual data points for that sample. Error bars reflect the standard error of the mean. T-tests were conducted to determine statistical significance (*p<0.05, **p<0.005, ***p<0.0005, ****p<0.00005).

GFP-positive cells did not vary for the readthrough control across all conditions, indicating that our control was working as expected (**Supplementary Fig. 4A-B**). There was a basal level of readthrough for both NMD+ reporters when treated with DMSO. This level significantly increased when treated with the readthrough-inducing drug G418 for the EJC-independent reporter but not for the EJC-enhanced reporter (**Fig. 4B** and **Supplementary Fig. 4A-B**). Upon SMG1i treatment, there was no increase in the percentage of GFP-positive cells for the EJC-independent reporters, however, there was a mild but significant increase for the EJC-enhanced reporters. When G418 and SMG1i were combined, there was an additive increase in GFP-positive cells for the EJC-independent reporter and a synergistic increase for the EJC-enhanced reporter (**Fig. 4B** and **Supplementary Fig. 4A-B**). Notably, these data show that the EJC-independent and EJC-enhanced reporters respond differently to the G418, SMG1i, and combined drug treatments.

GFP protein levels increased for the readthrough control upon treatment with G418 but not SMG1i. GFP protein levels increased to the same extent as with only G418 treatment upon combined drug treatment (**Supplementary Fig. 4C**). Once again, a basal level of GFP protein was detected for both NMD+ reporters when treated with DMSO (**Fig. 4C** and **Supplementary Fig. 4C**). These levels increased to similar levels when treated with G418 for both the EJC-independent and EJC-enhanced reporters. SMG1i treatment also increased GFP protein levels for both NMD+ reporters to similar levels. The increase was mild but significant and did not reach the GFP protein levels of reporters treated with G418. The combined treatment of G418 and SMG1i still induced a diverging response, with the EJC-independent reporter showing an additive increase and the EJC-enhanced reporter showing a synergistic increase in GFP protein (**Fig. 4C** and **Supplementary Fig. 4C**). Taken together, these data suggest that the NMD+ reporters are responsive to G418 and thus, can be used to study readthrough. However, the data cannot distinguish between direct translational readthrough or an increase in readthrough due to stabilization of the mRNA since these two processes are intricately linked. Nevertheless, given the differential responses of the NMD+ reporters to the drug treatments, these data highlight the importance of comparative studies between EJC-independent and EJC-enhanced NMD.

PTC readthrough is known to stabilize NMD targets, so we next looked at the RFP levels of each reporter construct across all treatments (**Fig. 4D**) [35]. The RFP protein levels of the readthrough control increased in all conditions that used G418, similar to the GFP protein levels for the same reporter. For the NMD+ reporters, G418 treatment caused a small but significant increase in levels of RFP, suggesting that both reporters are being stabilized. As expected, SMG1i treatment increased RFP levels of both NMD+ reporters. The combined drug treatment increased RFP to the same level as the cells with only SMG1i treatment for both reporters, thus, no additive or synergistic increase in RFP was observed corresponding to the level of readthrough. It is possible that the use of the TAG or TGA stop codons with G418 and SMG1i would lead to differing observations than the ones reported in this study, such as more potent readthrough, since these stop codons have lower fidelity compared to TAA [61–64]. The extent of readthrough, however, may still differ between EJC-independent and EJC-enhanced NMD.

## Conclusion

In this study, we introduce a reporter system that facilitates the investigation of multiple sources of variability on NMD, including substrate-type, and mode of expression in cells. We present 16 different reporter plasmids that include both fluorescent and luminescent versions of EJC-enhanced and EJC-independent NMD reporters as well as two different controls for each version of reporters, all of them provided with and without an intron in the 5’ UTR. Notably, our system integrates these sources of variability into a unified framework, enabling comprehensive analysis within a single experimental setup. It also has the potential to identify and detect variability in NMD-associated phenomena like PTC readthrough and truncated protein degradation. Furthermore, it is compatible with high throughput assays such as genetic screens and massively parallel reporter assays. Consequently, it has the potential to advance our understanding of multiple aspects of NMD biology and regulation. These reporters are available to the scientific community through Addgene.

Existing fluorescence- and luminescence-based reporters (**Figure 5**) have enabled important discoveries pertaining to the NMD pathway including novel modulators of NMD, the variability of NMD efficiency within and between cell types, and the study of NMD-associated phenomena such as PTC readthrough and truncated protein degradation. The reporters developed in this study fulfill an important need to test our current understanding of NMD comparatively between the two modes of NMD to yield insights into the nuances of NMD and factors that might affect the two pathways differentially. To allow the NMD community to take advantage of the entire reporter system landscape available, we have compiled a table delineating the features of different reporters (including the ones developed in this study) and the types of investigations for which they are suitable (**Figure 5**).

**Fig. 5:**
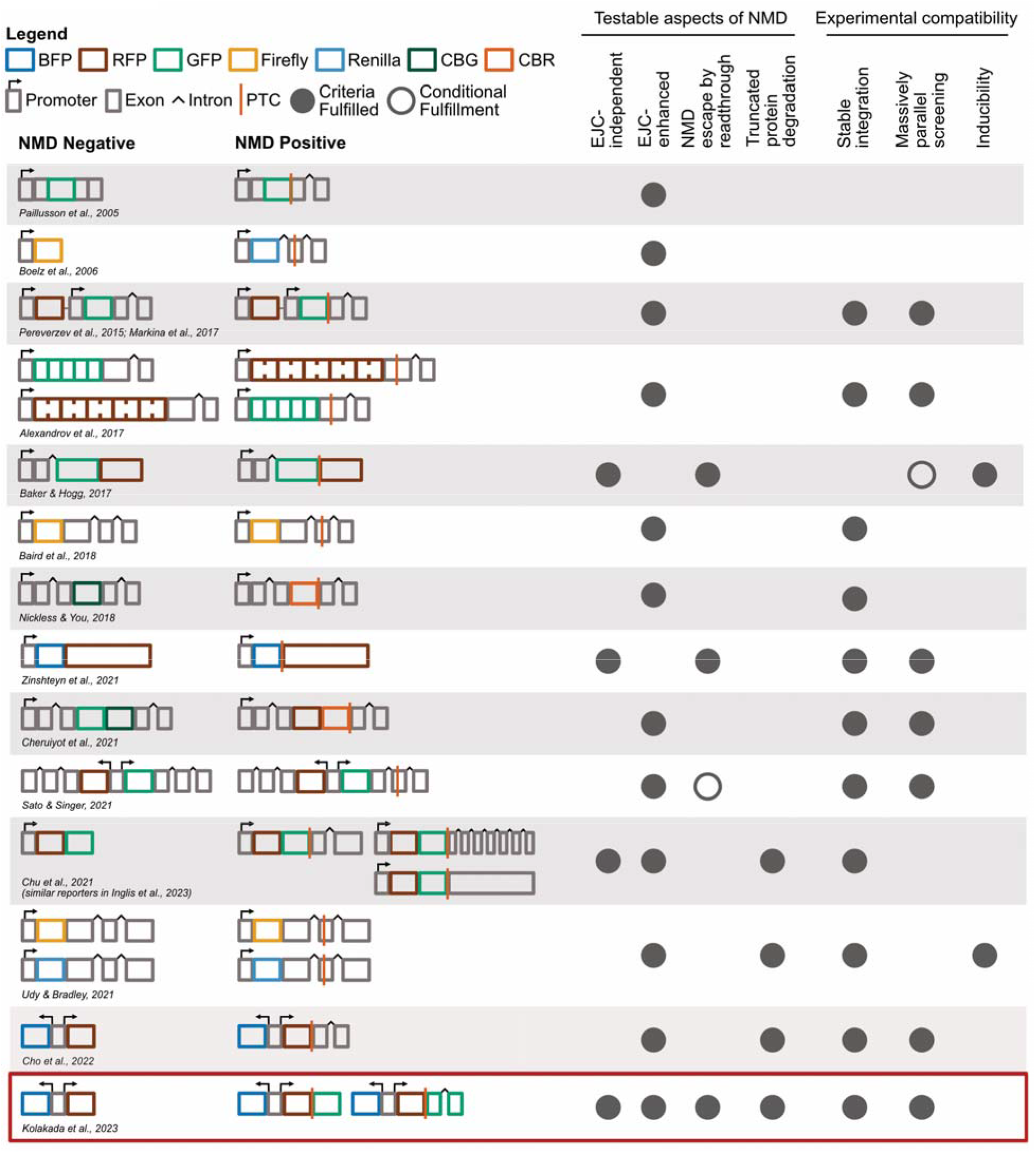
The NMD reporter landscape. A summary of existing NMD reporter systems, their ability to test various aspects of NMD and their experimental compatibilities. Each reporter system includes the NMD+ reporter and its NMD– control. Fluorescent or luminescent reporter genes are represented by colored boxes (CBG: Click Beetle Green luciferase, CBR: Click Beetle Red luciferase), gray boxes represent exons of known NMD targets, and black carets indicate their introns. Various promoter elements driving reporter gene expression are shown as gray boxes with arrows, and orange lines denote PTCs. Testable aspects of NMD includes to the two branches of NMD and the NMD associate mechanisms PTC readthrough and truncated protein degradation. Experimental compatibilities cover stable recombinase-mediated genomic integration, massively parallel reporter assays, and inducibility if the reporter has an inducible promoter. Filled gray circles indicate fulfillment of a criterion, while empty gray circles indicate conditional fulfillment. The *Sato and Singer* reporters are conditional for readthrough because GFP protein levels cannot distinguish between readthrough and truncated protein expression. The *Baker and Hogg* reporters are only suitable for massively parallel screening if they can be stably integrated. The reporters introduced in this study are highlighted with a red box.

## Data Availability

The data underlying this article are available in the article and in its online supplementary material.

## Acknowledgements

We thank Heather Johns, Qing Feng, and other members of Dr. Robert Bradley’s lab for feedback on early versions of these reporter constructs. We thank members of the Jagannathan Lab for insightful manuscript feedback. We thank the Taliaferro Lab for providing us with the RMCE donor plasmid and Hek293T LoxP cell lines.

