## Supplementary figures for "A system of reporters for comparative investigation of EJC-independent and EJC-enhanced nonsense-mediated mRNA decay"

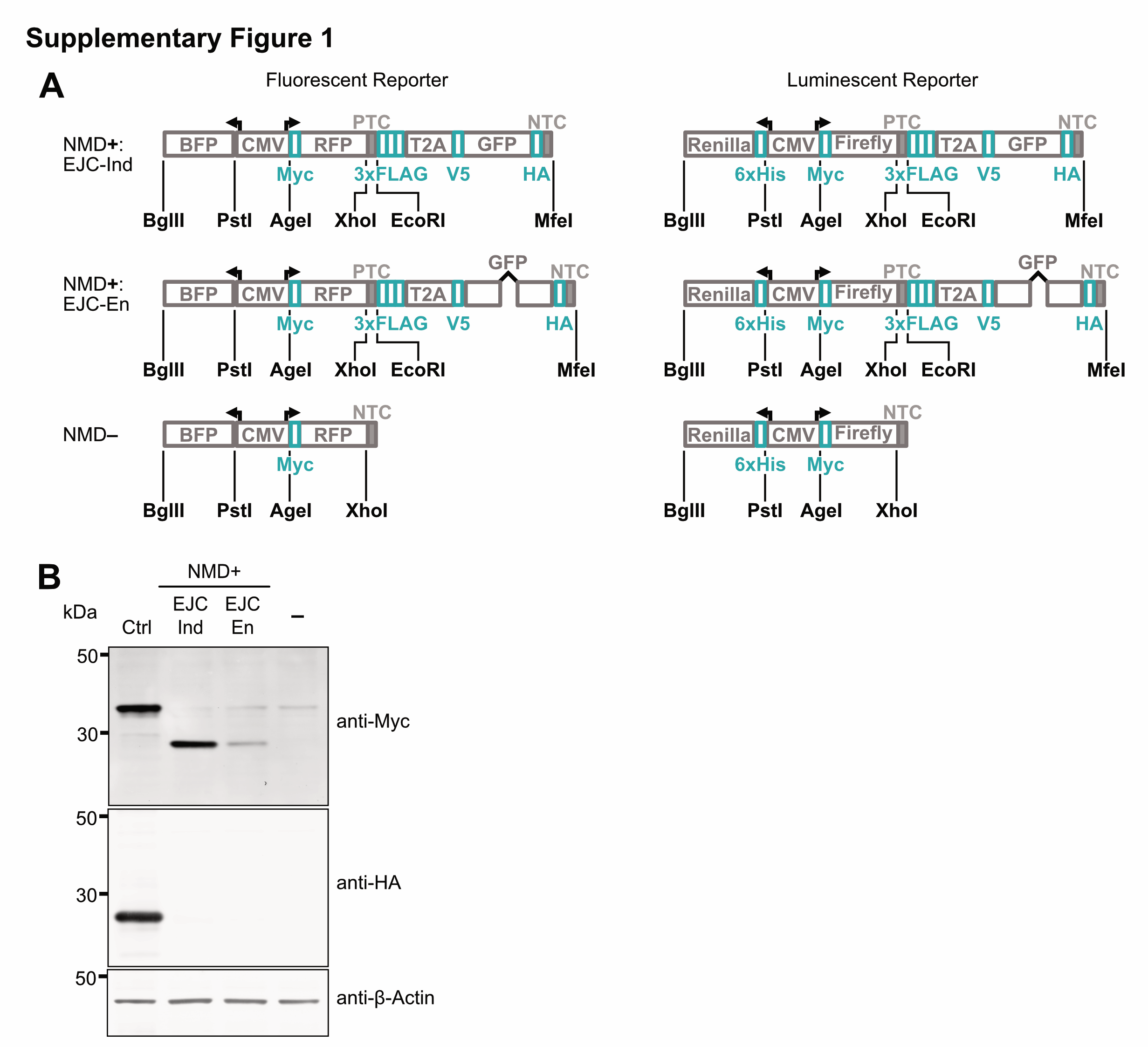

**Supplementary Fig. 1:** (A) A detailed diagram of both the fluorescent and luminescent NMD reporters showing all restriction sites available to modify the reporters (black) as well as the tags on each reporter protein to enable western blotting or pull-down (teal). (B) Western blot to detect RFP and GFP with N- and C-terminal tags. Top panel: Anti-myc which is fused to the N-terminus of RFP. Middle panel: Anti-HA which is fused to the C-terminus of GFP. β-Actin was used as the loading control. Ctrl refers to a construct that lacks the stop codon between RFP and GFP, to enable visualization of both proteins. EJC Ind and EJC En refer to the EJC-independent and EJC-enhanced NMD+ fluorescent reporters, respectively. The “–” sample refers to a no transfection control. Samples were generated from HEK293 cells harvested 48 hours post-transfection.

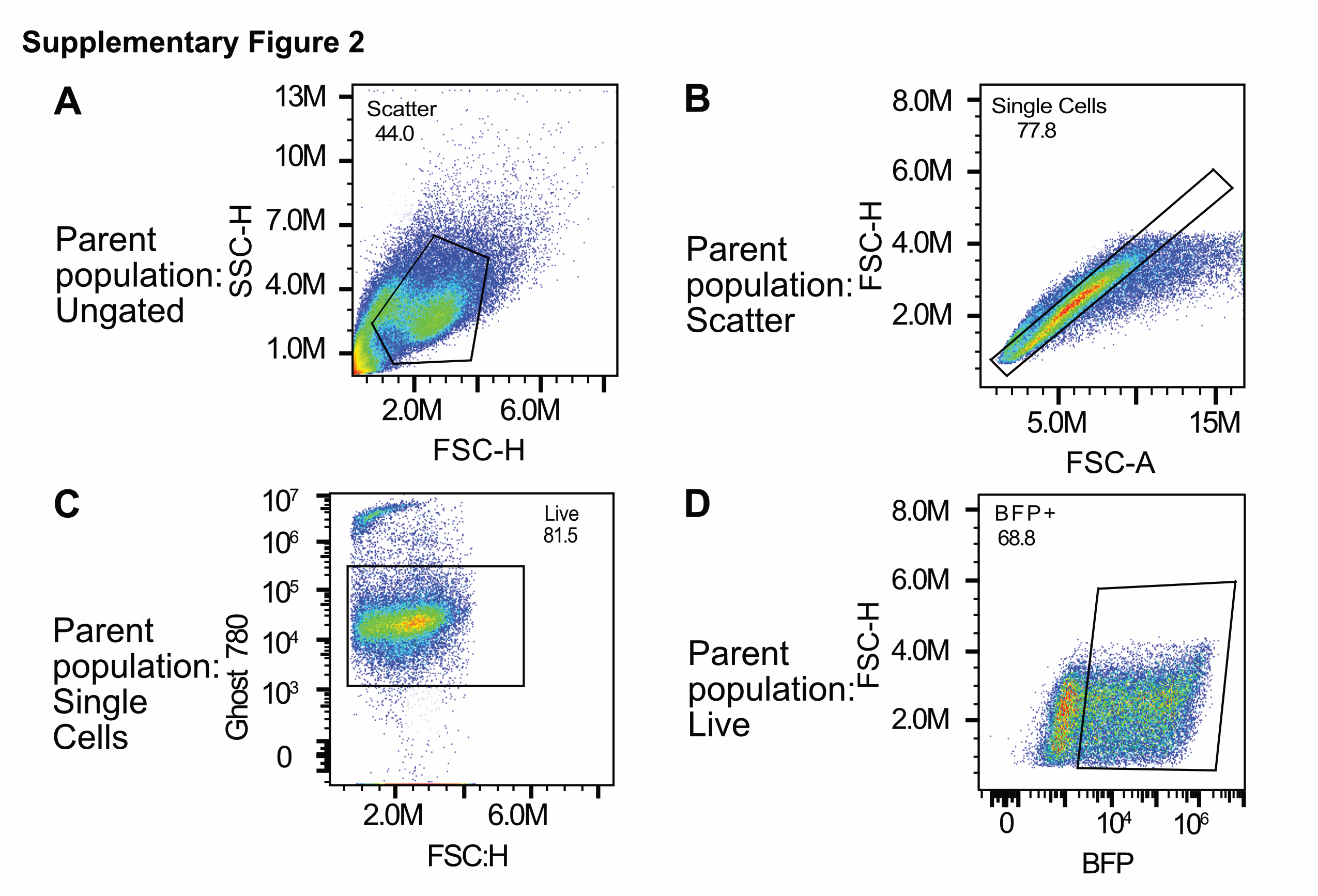

**Supplementary Fig. 2:** Gating strategy for flow cytometry. (A) Size-based selection of cells of interest from the ungated cell population. (B) Exclusion of doublets and selection of only single cells from the scatter population. (C) Selection of live cells via staining with Ghost 780 viability dye from the single cell population. (D) Selection of BFP+ expressing cells from live cells.

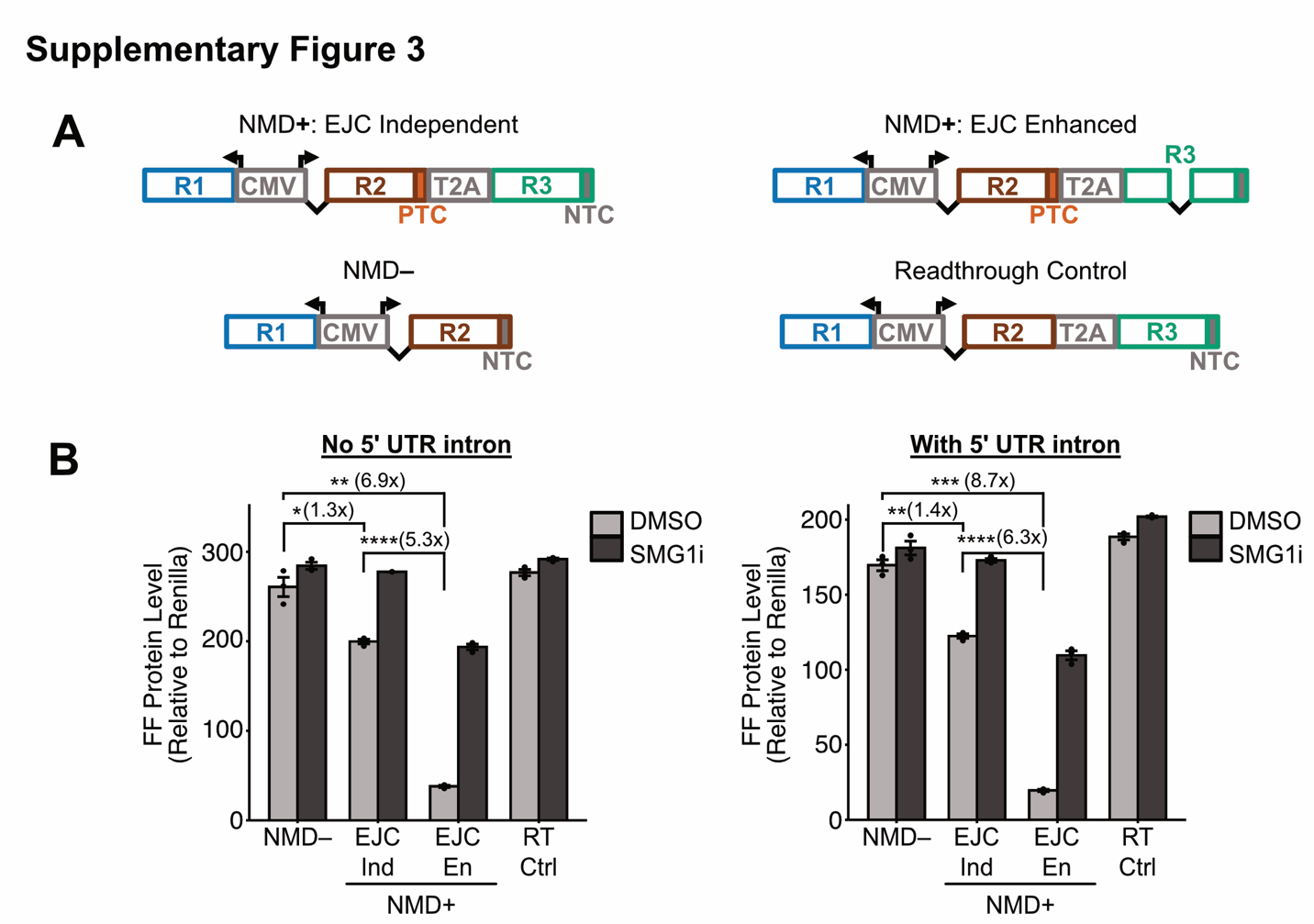

**Supplementary Fig. 3:** Inclusion of a 5’ UTR intron does not alter the relative behavior of the NMD reporters. (A) The NMD+: EJC-independent, NMD+: EJC-enhanced and NMD– reporter and readthrough control designs including a synthetic intron in the 5’ UTR. The thin black line connecting exon blocks refers to the intron. (B) Protein levels of luminescent NMD reporters with and without a 5’ UTR intron. Left: Firefly (FF) luciferase protein levels of the original NMD– and NMD+ vectors with no intron in the 5’ UTR. EJC Ind refers to EJC-independent reporter, EJC En refers to the EJC-enhanced reporter, and RT Ctrl is the luminescent readthrough control with firefly luciferase and GFP in a single open reading frame with no stop in between. Right: Same plot as at left, except using reporters containing an intron in the 5’ UTR. For both experiments, transfections were performed in triplicates and the mean values of protein levels were plotted in the bar graphs. Dots overlaid on each bar indicate the individual data points for that sample. Error bars reflect the standard error of the mean. T-tests were conducted to determine statistical significance (*p<0.05, **p<0.005, ***p<0.0005, ****p<0.00005).

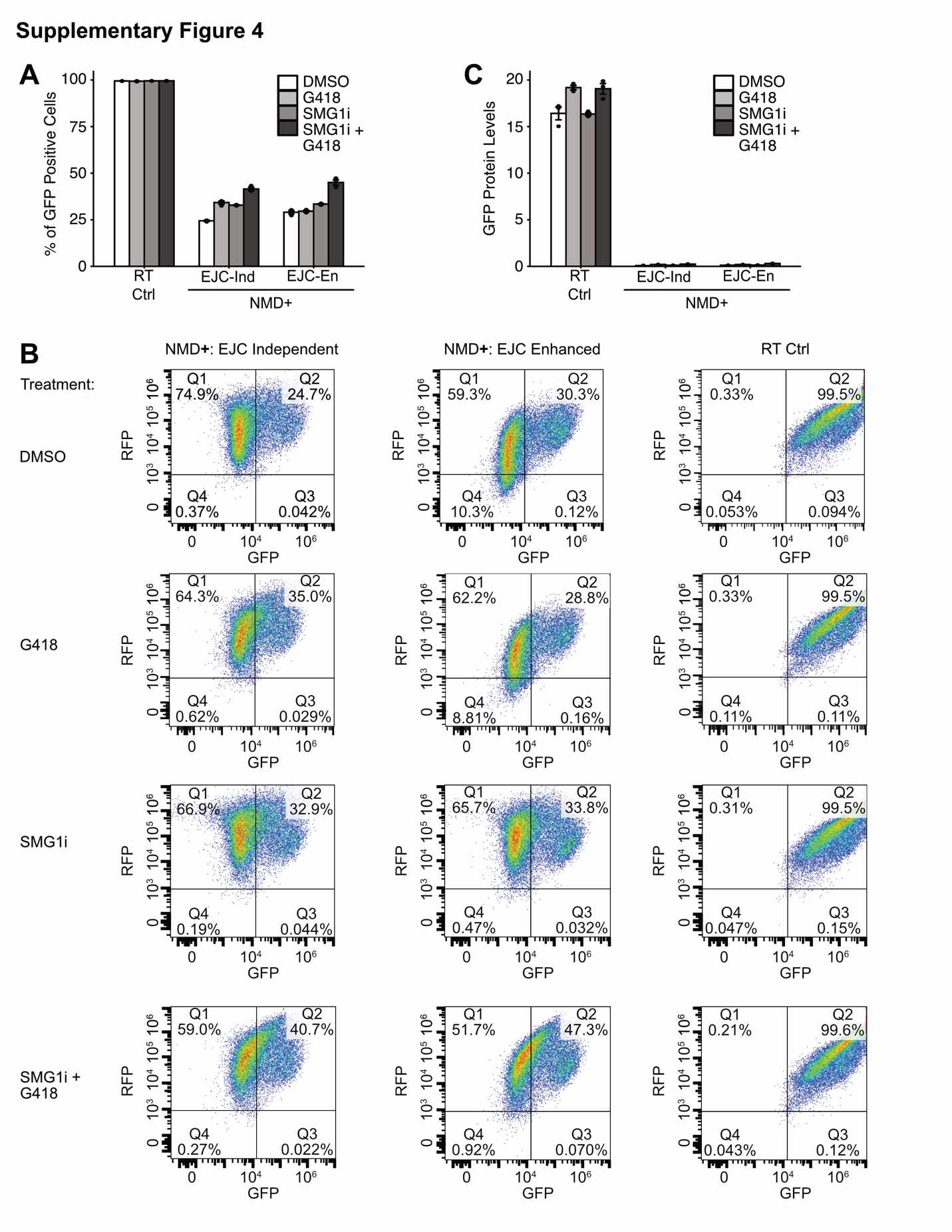
**Supplementary Fig. 4:** Percentage of GFP+ cells does not vary for the readthrough control across all conditions. (A) Percentage of GFP positive cells for all reporters when treated with DMSO, G418, SMG1i, and SMG1i+G418. (B) Representative flow diagrams across all reporters and treatments, the percentage of GFP positive cells are in Q2. (C) Mean GFP protein level for all reporters when treated with DMSO, G418, SMG1i, and SMG1i+G418. For all experiments, transfections were performed in triplicates and the mean values of protein levels were plotted in the bar graphs. Dots overlaid on each bar indicate the individual data points for that sample. Error bars reflect the standard error of the mean. T-tests were conducted to determine statistical significance (*p<0.05, **p<0.005, ***p<0.0005, ****p<0.00005).
